# Electrical stimulation of grape berries as new alternative for post-harvest infection management

**DOI:** 10.1101/2024.04.15.589490

**Authors:** Mariana Fernandes, Flávio Rodrigues, Vanessa F. Cardoso, Filipe Silva, Margarida M. Fernandes

**Affiliations:** Center for MicroElectromechanical Systems (CMEMS-UMinho), University of Minho, Campus de Azurém, 4800-058, Guimarães, Portugal; LABBELS-Associate Laboratory in Biotechnology and Bioengineering and Microelectromechanical Systems, Universidade do Minho, Braga/Guimarães, Portugal

**Keywords:** grapes, electrical stimuli, infection, *Escherichia coli*, pest management

## Abstract

Viticulture stands out as a primary agricultural sector in the EU. However, an intensive regime of chemical pesticides is used to meet production standards. Consequently, the EU has taken measures to encourage the promotion of sustainable agrochemical practices. It has recently been demonstrated that low voltage and electrical current can also successfully destroy different type of microorganisms such as bacteria and fungi. However, the application of electric current has not been much explored for preventing early stages of fresh fruit rotting. As a result, an innovative setup was developed to study the effect of an alternating current (50 mA) on grapes previously contaminated with *E. coli*. A visual analysis of the state of degradation of the grapes 12 days after the electrical treatment allowed us to conclude that the contaminated grapes subjected to electrical treatment (at day 1) did not show visual contamination, when compared to the contaminated grapes without electrical treatment, presenting a similar appearance to the grapes used as control. The quantitative assay also confirmed this result since on the NB agar plates it was possible to visualize a significant reduction in colonies, corresponding to 2 log_10_ CFU/mL, indicating that approximately 99% of the *E. coli* previously attached to the berry’ surface was eradicated. This environmentally sustainable solution aims to reduce dependence on pesticides. Still, other studies will focus on validating its effectiveness against target microorganisms and optimizing a conductive network for applying electrical stimuli to a bunch of grapes.

## 1. Introduction

The European Union (EU) is the world leader in grapevine production, with approximately 3.2 million hectares, which represents 45% of the total vineyard area worldwide. Despite its socioeconomical importance, viticulture confronts huge challenges since it mainly relies on agrochemicals to prevent vines’ susceptibility to fungal and bacterial infections. *Plasmopara viticola, Botrytis cinerea, Aspergillus niger, Penicillium expansum, Erysiphe necator* and *Esca*-associated pathogens, as well as bacteria like *Rhizobium* (*Agrobacterium*) *vitis* are common pathogens in the EU’s grapevines, being mainly present in the grape berry’s surface. Climate change is increasing the prevalence of these pathogens, which is having a severe detrimental impact on grape production, including reduced yields and degraded quality (Di Canito et al., 2021; Eastwell et al., 2006; Karácsony et al., 2023; Pennington et al., 2018; Pertot et al., 2017).

Control of pests is often managed with the use of pesticides. However, combined with excessive costs, they may have unexpected repercussions on human health and environment due to their high biological activity and persistence, which is forcing EU (Directive 2009/128/EC) to ban many synthetic pesticides, establish rules for sustainable use of agrochemicals and explore new alternatives. These approaches refer to an arsenal of new, non-chemical solutions, to combat fruit and vegetable diseases, promoting food production and conservation (D’Addabbo et al., 2014; Di Canito et al., 2021; Maroni et al., 1999). A promising solution involves using physical agents, as potential substitutes for antibiotics (Marian Darma et al., 2022).

In recent years, decontamination through the application of physical agents, such as ultrasound, to fresh fruits and vegetables has been widely studied for application in the food industry (São José et al., 2014). Piezoelectric materials have been also explored, which create electroactive microenvironments at their surface when mechanically stimulated (Carvalho et al., 2019; Costa et al., 2023; Moreira et al., 2022). It has also been shown that magnetoelectric materials induce bactericidal effects when magnetically stimulated (Fernandes et al., 2021; Marqués-Marchán et al., 2023a). Unfortunately, there are still far too few of these experiments based on the use of smart and functional materials for the creation and improvement of new techniques. This might be related to the difficulty in standardizing methods as well as the multiple aspects that must be considered at the same time (voltage or current intensity, eventual usage of electrodes, and duration of treatment) (Krishnamurthi et al., 2020; Ranalli et al., 2002; Valle et al., 2007).

Other interesting physical stimulus is the application of electric current that have been studied as an alternative to chemical disinfectants for inhibiting and/or killing microorganisms since the 1960s. Electric current is currently targeted for the development and optimization of revolutionary approaches for water and food disinfection and/or conservation, although its application is still in its early stages. The primary goal of such methods is to diminish or remove some of the frequently unwanted microorganisms present (Krishnamurthi et al., 2020; Ranalli et al., 2002; Valle et al., 2007).

Some studies have reported the effects of low electric current on yeast cells. Ranalli *et al*. found that 10, 30 and 100 mA, with the electrode positioned at the bottom of the well, could reduce the adenosine triphosphate (ATP) content and the viability of suspension cultures of *S. cerevisiae* and *H. guilliermondii*. Furthermore, the low electric current treatment of *S. cerevisiae* resulted in cytoplasmatic membrane integrity degradation (Ranalli et al., 2002). Another study showed that when applying 200 or 2,000 μA for 4 days, to electrodes inserted 3 mm from discs with pre-formed biofilms, several strains of *C. glabrata* and *C. albicans*, respectively, showed a significant reduction in biofilm (Schmidt-Malan et al., 2015).

Different tests were carried out on bacterial biofilms (*S. aureus, S. epidermidis, P. aeruginosa*) adhered to discs with electrodes positioned on both sides, 3 mm from the disc, using 200 μA. The results showed a reduction in microbial viability due to changes in cell morphology. Furthermore, reactive oxygen species (ROS) appear to play a role in cell death associated with direct current application, as there was an increase in ROS production by *S. aureus* and *S. epidermidis* biofilms following exposure to direct current (Brinkman et al., 2016). Valle *et al*. also showed that there was a notable inhibition of enzyme activities, growth and a reduction in the ATP content of the *E. coli* bacterial suspension using 20 and 40 mA. This test was carried out in cylindrical glass tanks (20 cm in diameter, 10 cm in height) and the electrode pairs were placed in the tanks at a distance of 19 cm from each other (Valle et al., 2007).

In this work, an innovative setup was developed to study the effect of an alternating current (50 mA) on grapes contaminated with *E. coli*. The effects of the low electric current intensity on bacterial viability were evaluated using conventional cultural method (viable cells counts) and visual assessment. This work demonstrates the feasibility of using electric stimuli on fruit to mitigate early rotting caused by microbial contamination.

## 2. Materials and methods

### 2.1 Microorganisms and media

Pure cultures of *E. coli* (CIP 110067) were used. Laboratory bacterial assays were performed in Tryptic Soy Broth (TSB) medium (Merck, Darmastadt, Germany). The bacteria were first grown at 37 °C overnight with orbital rotation at 250 rpm. On the second day, the pre-inoculum was centrifuged at 5000 rpm for 5 min and washed twice with fresh medium. The optical density was measured at 600 nm (OD_600_) and adjust to 0.1 ± 0.01 using TSB medium. This adjustment resulted in a working inoculum of approximately 7.3 × 10^7^ CFU mL ^-1^.

### 2.2 Grapes Washing and *E. coli* Contamination

The grapes used in this study were purchased from a local supermarket. To avoid interference with their own microbiota in the tests with *E. coli*, the grapes were first washed in a cleaning solution containing 0.05% (v/v) sodium hypochlorite (bleach) for 1 h at 500 rpm and then washed with plenty of water to remove excess of bleach. For the infection process with *E. coli*, a sterile 12-well plate was used, in which a grape berry was placed in each well containing an *E. coli* solution at OD_600_ of 0.1, and incubated overnight, i.e. approximately 12 h, at 37 °C.

### 2.3 Electrical stimulation actuator development

To electrically stimulate the grape berry, an electrical setup was built for delivering a well-defined electrical current to the grapes, using NaCl 0.9% as electrolyte. The setup, schematically represented in **Figure 1**, consists of two Teflon discs positioned above and below a network of 12 silver wires (0.3 mm diameter), which serve as electric conductors. Placed between the discs and inside the network of wires are the grapes, serving as the study elements. Within the network, consecutive wires act as opposite electrodes (6 anodes and cathodes, alternately actuated), forming a repeating pattern around the grapes (**Figure 1**). An electric insulator was placed in contact points to guarantee that wires with opposite charges are not in contact. This setup was built to create several sites for current flow sites and form a network around the whole grape berry. Teflon discs were chosen to construct the setup apparatus due to their biological inertness (Korzinskas et al., 2018; Schmidt-Malan et al., 2015). On the other hand, silver wires were used to conduct the electrical current since it is an inorganic material renowned for its antimicrobial properties and high electrical conductivity (Chouirfa et al., 2019; HOLLER et al., 2007).

**Figure 1.**
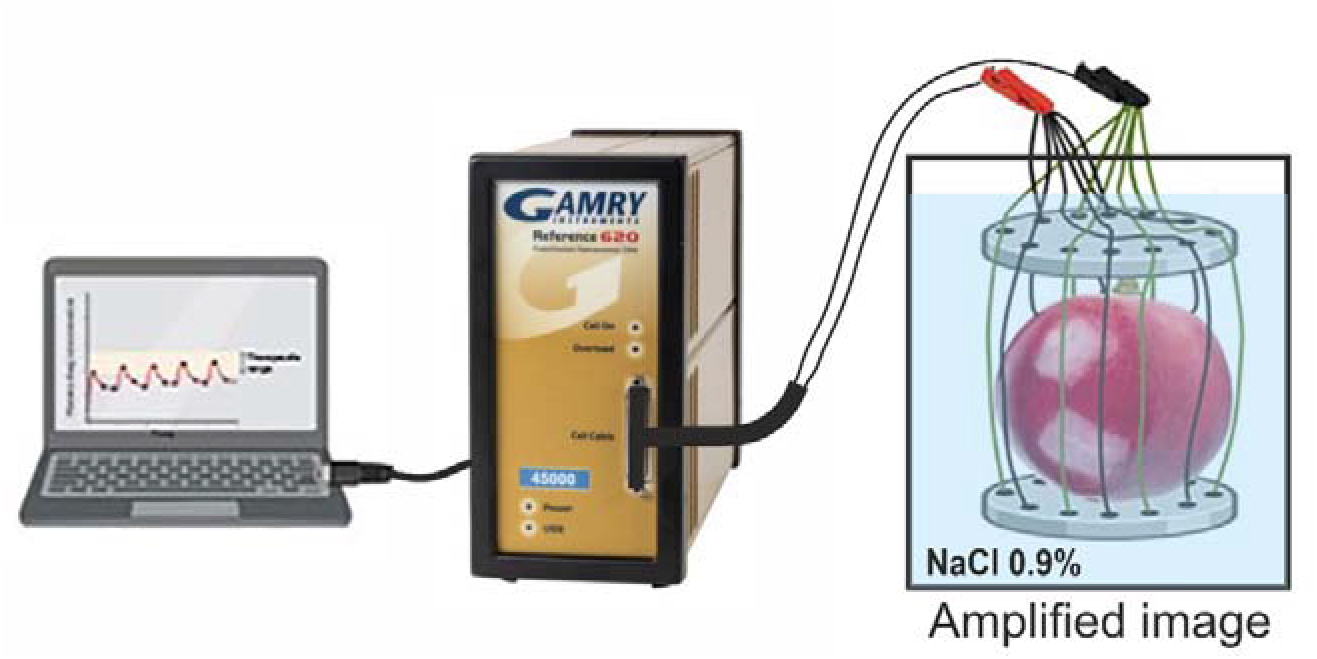
Experimental setup designed for the electricity study. The green wires represent the positive pole and the black wires the negative pole immersed in NaCl 0.9% as electrolyte.

### 2.4 *In vitro* Electrical stimulation of grapes

After overnight infection with *E. coli*, the grape berries were then individually treated in a well of a sterile 12-well plates (0.025 cm^2^ area) through the application of a defined electrical current using a current generator (Gamry®600+). The Gamry®600+ was programmed to deliver a constant current of 50 mA with polarity inversion every 1 s for 10 min (300 cycles) while monitoring the potential variation. This means that the electric current (*i*.*e*., electrons) enters through the negatively charged electrodes (cathodes); solution components travel to these electrodes, combine with the electrons, and are transformed (reduced), cyclically. This electrical density was chosen following previous studies on the application of electrical current, which have shown that current as low as 25 mA has the capacity to kill the *E. coli* bacteria (Pareilleux & Sicard, 1970). Teflon disks and the electrode network device were sterilized before and after each assay by subjecting them to ultraviolet (UV) light for 30 min. For the experiments, each contaminated grape was placed inside and in contact with the electrode network of the actuator and submerged in 3 mL of a sterile NaCl 0.9% (w/v). Two control tests were prepared: i) non-contaminated grapes (only subjected to initial washing) and ii) contaminated grapes and not subjected to current application. All tested grapes, including controls, were divided into two groups for: visual assessment over time and quantitative assessment using the Colony-forming-unit (CFU) assay. Six replicates were performed for each sample.

### 2.5 Qualitative and quantitative studies on the effect of electrical current on contaminated grapes

For a qualitative analysis, all grapes were immediately placed inside a sterile zipper bag and their contamination/degradation was visually analysed over time. In a quantitative study, the grapes that were subjected to current application were immediately washed in a 3 mL of a NaCl sterile solution (0.9% (w/v)) for 30 min at 500 rpm. The two controls were submerged for 10 min in a 0.9% (w/v) NaCl sterile solution (to remain in the same conditions as the stimulated grapes) and subsequently washed in 3 ml of a NaCl sterile solution (0.9% (w/v)) for 30 min at 500 rpm. The suspensions obtained were subjected to 8 serial dilutions (1:10) in a sterile 0.9% (w/v) NaCl solution. For each dilution, a volume of 20 μL was dropped on Nutrient Broth (NB, Merck, Darmastadt, Germany) plate agar and further incubated at 37 °C for 24 h. Following, the number of viable bacteria was determined, allowing a quantitative analysis of the average CFU⋅mL^− 1^. The bactericidal activity was evaluated with three independent assays as log_10_ for each condition.

### 2.6 Statistical Analysis

The results were statistically analyzed, and the average of individual measurements with the respective standard deviations was used. The significance of differences between contaminated grapes and electrically treated grapes treatments was assessed using Student’s *t*-test. The tests were performed using the software GraphPad Prism version X for Windows (GraphPad Software, San Diego, CA, U.S. A.)

## 3. Results and discussion

Individual grape berries were electrically stimulated to show the potential of using alternative non-chemical ways for preventing microbial-induced rotting in grapes. For that, an actuator was built to allow the application of an electrical current on the surface of each grape berry individually (**Figure 1**). It was important to assure that the grapes skin was intact and no leakage from the interior was observed, as the interior of the grape may also bey colonized by their own microbiota.

**Figure 2** illustrates a 100-second segment of the aforementioned experiment. Throughout the process, the alternating current remained effectively constant (± 50 mA). However, according to the experimental setup designed for the electricity study (**Figure 1**), it can be measured, despite the applied current being 50 mA, for each of the 12 wires that surround the grape, the current is divided and, therefore, theoretically for each wire will pass about 4.17 mA.

**Figure 2.**
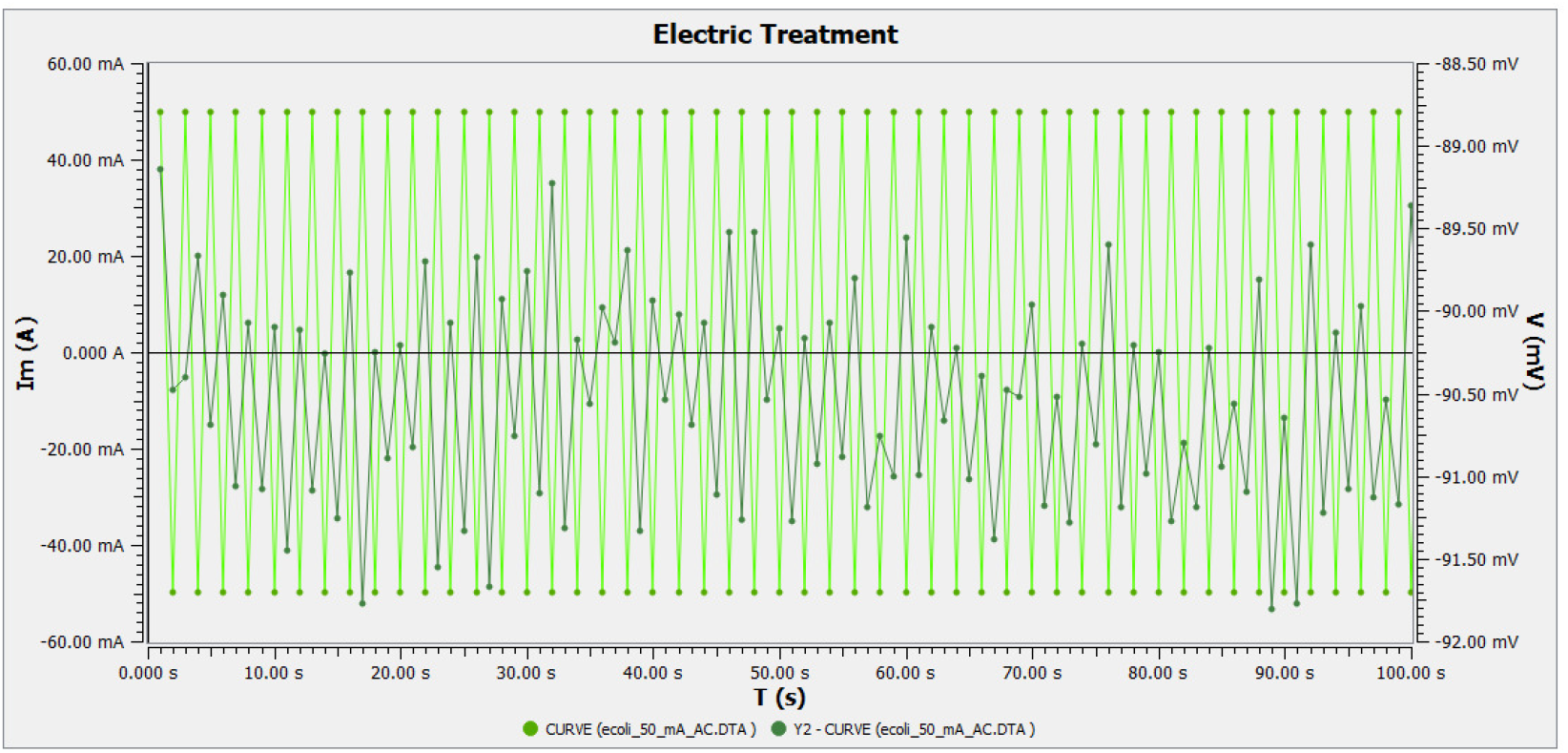
100-second segment of the electrical treatment acquired by Gamry software: current (A) and potential (mV).

Regarding the potential difference, there were variations in potential, with maximum and minimum values recorded at -91.93 mV and -88.87 mV, respectively. The device included a feature to adjust the potential to maintain a constant electric current, which was the variable under control. The observed variations in potential could be attributed to fluctuations in resistance within the circuit. According to Ohm’s Law (V = I * R), if the current (I) remains constant while the potential (V) varies, it indicates a change in resistance (R) within the circuit (Carter, 2013). The presence of adherent cells in the grape, and in the medium, can affect the current flow by increasing or decreasing the resistance of the circuit. The magnitude of this resistance depends on the number of bacteria and their size, and can also be associated with metabolic activity and the extracellular polymeric substance that acts as an insulating layer (Gutiérrez et al., 2016; Paredes et al., 2014).

As previously stated, the contamination of the grapes was carried out in a controlled manner, contaminating all the grapes with the same amount of bacteria (OD_600_=0.1) and for the same time, i.e. approximately 12 h, as a way to guarantee effective and dispersed contamination on the surface of all grapes. Then, individual grape berries were electrically stimulated under controlled current conditions, previously described, to show the potential of using alternative non-chemical ways for preventing microbial-induced rotting in grapes. It was important to assure that each grape was totally immersed in the electrolyte NaCl 0.9% to guarantee homogeneous electrolysis at the surface of the grape. Moreover, the skin should remain intact and no leaks must be observed from the inside, as the interior of the grape may also bey colonized by their own microbiota. Thus, the antibacterial tests were performed on contaminated grape berries to evaluate the capacity of low electrical current to eradicate the clinically relevant Gram-negative bacteria *E. coli* previously attached to the berry’s surface. The results were then compared to the controls. The bactericidal activity of the application of milliampere current was evaluated against the clinically relevant Gram-negative bacteria *E. coli*. This bacterium was used as a representative model of a Gram-negative bacterium, to mimic one of the most pathogenic bacteria in vines, the *Rhizobium* (*Agrobacterium*) *vitis*, a bacterium that causes gall disease and is also a Gram-negative bacterium. Furthermore, *E. coli* is already considered one of the six main killer pathogens associated with antimicrobial resistance (Eastwell et al., 2006; Marqués-Marchán et al., 2023).

According to the visual analysis of the state of contamination/degradation of the grapes (**Figure 3A**), it was found that after 12 days, the contaminated grapes subjected to electrical treatment (at day 1) did not show visual contamination, when compared to the contaminated grapes without electrical treatment, presenting a similar appearance to the grapes used as control (washed and microorganisms-free grapes). These preliminarily results indicate that both the washing of the grapes and the contamination were done effectively.

**Figure 3.**
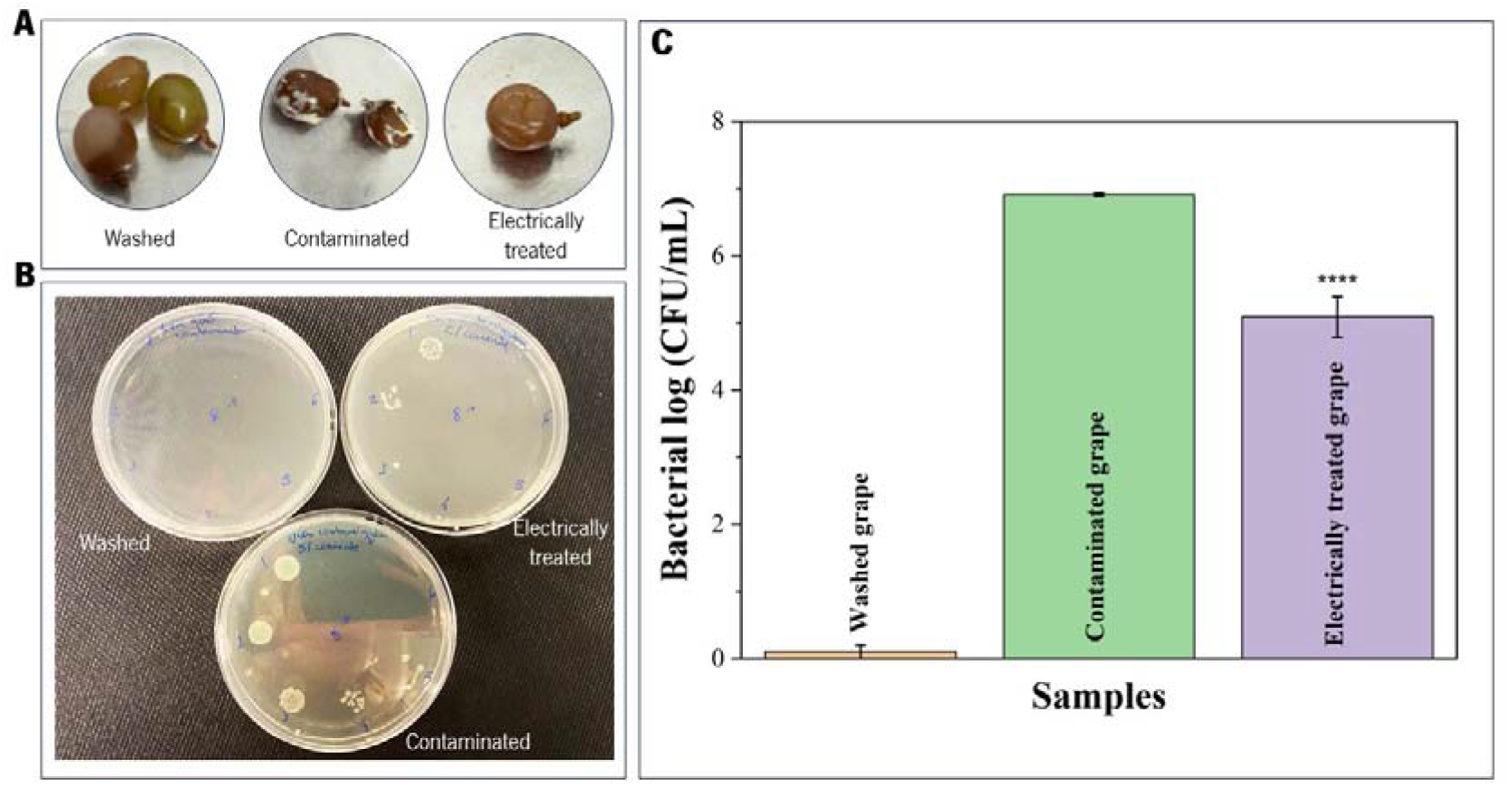
Evaluation of the bactericidal activity of grapes: wash, contaminated by the *E. coli* bacteria without current treatment and contaminated by the *E. coli* bacteria after applying 50 mA current for 10 min (at day 1): **A)** Visual appearance 12 days after starting the test; **B)** Quantitative study based on CFU; **C)** Bacterial activity as log_10_ (CFU/mL) of the suspensions obtained after the test with grapes. ****P < 0.0001 when compared contaminated grapes with electrically treated grapes.

In fact, the quantitative test confirmed the previous assumption, since there was no growth of microorganisms in the respective washing control and, in the contaminated control, only colonies from *E. coli* strain were observed (**Figure 3B**). When comparing contaminated grapes, it can be observed that the application of 50 mA for only 10 min at day 1 induced a significant reduction in bacterial cells at the grape’ surface (electrically treated grapes). In fact, for the same dilution, the number of colonies presented on the test plate agar, corresponding to the electrical treatment, is significantly lower than the number of colonies visible on the agar plate corresponding to the contaminated control (**Figure 3B**). After application of the electrical stimulus, the reduction was approximately 2 log_10_ CFU/mL, (**Figure 3C**), indicating that approximately 99% of the *E. coli* previously attached to the berry’ surface was eradicated. This difference in bacterial reduction is statistically significant (****P < 0.0001), highlighting the bactericidal effect capacity of electrical stimulus.

Although the predominant pathogens in EU grapevines are fungi and *Agrobacterium*, similar effects are expected in the main fungal pathogens, if we considered that electric currents can disrupt the cell membrane of both bacterial and fungal cells, leading to cell disruption that can lead to leakage of cellular contents, loss of cell integrity, and ultimately cell death. In fact, the lethal action of electric fields is governed mainly by the intrinsic properties of the microorganism, under which the composition of the cell membrane may play the most important role (Hülsheger et al., 1983). Furthermore, the lethal activity of milliampere and microampere current in yeasts has already been confirmed by other reports (Ranalli et al., 2002; Schmidt-Malan et al., 2015).

## 4. Conclusions and future perspectives

This preliminary study was performed to prove the concept of electrical current in preventing grape rotting. In fact, a decrease or even 99% eradication of *E. coli* in the grapes was achieved using electrical currents as low as 50 mA for just 10 min. The overarching aim is to extend the applicability of the developed methodology to target other microorganisms, particularly prevalent pathogens affecting grapevines within the EU region. This proposed solution represents an environmentally sustainable approach that mitigates reliance on conventional pesticides and chemical agents. Further studies will focus on the validation of this protocol against the specified microorganisms of interest and the development and optimization of a polymeric conductive net that allows for the application of the electrical stimuli in a bunch of grapes.

## Declaration of Competing Interest

The authors declare that they have no known competing financial interests or personal relationships that could have appeared to influence the work reported in this paper.

## Acknowledgments

The authors thank the FCT-Fundação para a Ciência e Tecnologia for financial support in the framework of the Strategic Funding UIDB/04436/2020, UIDP/04436/2020, and the project 2022.02697.PTDC. VFC and MMF thanks the FCT for the contracts under the Stimulus of Scientific Employment, 2020.02304.CEECIND and CEECINST/00018/2021, respectively.

## Notes

### Competing Interest Statement

The authors have declared no competing interest.

